# Evaluation of lamin A/C mechanotransduction under different surface topography in LMNA related muscular dystrophy

**DOI:** 10.1101/2022.01.03.474777

**Authors:** Subarna Dutta, T. Muraganadan, Madavan Vasudevan

## Abstract

Most of the single point mutations of the LMNA gene are associated with distinct muscular dystrophies, marked by heterogenous phenotypes but primarily the loss and symmetric weakness of skeletal muscle tissue. The molecular mechanism and phenotype-genotype relationships in these muscular dystrophies are poorly understood. An effort has been here to delineating the adaptation of mechanical inputs into biological response by mutant cells of lamin A associated muscular dystrophy. In this study we implement engineered smooth and pattern surfaces of particular young modulus to mimic muscle physiological range. Using fluorescence and atomic force microscopy we present distinct architecture of the actin filament along with abnormally distorted cell and nuclear shape in mutants, which showed a tendency to deviate from wild type cells. Topographic features of pattern surface antagonizes the binding of the cell with it. Correspondingly, from the analysis of genome wide expression data in wild type and mutant cells, we report differential expression of the gene products of the structural components of cell adhesion as well as LINC (linkers of nucleoskeleton and cytoskeleton) protein complexes. This study also reveals mis expressed downstream signaling processes in mutant cells, which could potentially lead to onset of the disease upon the application of engineered materials to substitute the role of conventional cues in instilling cellular behaviors in muscular dystrophies. Collectively, these data support the notion that lamin A is essential for proper cellular mechanotransduction from extracellular environment to the genome and impairment of the muscle cell differentiation in the pathogenic mechanism for lamin A associated muscular dystrophy.

## Introduction

The microenvironment (for example, extracellular matrix mechanics) that surrounds the cell plays an important role in cell behaviour. Extracellular matrix stiffness (Janmey P et al. 2007,2011), deposition (Dai LY et al. 2011), topography (Spatz JP et al. 2004), migration and spreading through constriction (Saif MTA et al. 2014, Wang YL et al. 2005, Wolf KJ et al. 2010, Wang YL et al. 2000, Griffith LG et al. 2011,) ( Herman IM et al. 2016, Vogel V et al. 2014, Wong JYJ et al. 2008, Kemkemer R et al. 2010) are the determinants for the cellular mechanical homeostasis. In particular, matrix-based cues as well as cell-matrix interactions influence the disease aetiology in order to convert mechanical stimuli to biochemical signals, a process known as mechanotransduction. Integrin molecules mediate the first interaction between ECM and cell. Then, with the assistance of focal adhesion proteins, it travels to the cytoskeleton. It is then transferred to the nucleus by cytoskeleton elements via LINC complexes. Finally, this mechanotransduction cascade affects gene expression homeostasis. The mechanotransduction cascade is tissue specific. Mechanical characterization of tissues is heavily influenced by its implications for various biomechanical forces such as matrix stiffness. Biomedical engineering applications were used to determine the physiological ranges of various tissues. Optimal physiological ranges of tissues were determined using viscoelastic biopolymers and complex feedback systems, ranging from 100 Pa (in neural tissue) to 10kPa (in muscle tissue) and 1GPA (in bone) (Fratzl P et al. 2006, Janmey PA et al. 2002, Patrick CWJ et al. 2005, Aebi U et al. 2009, Van Vilet KJ et al. According to previous reports, young modulas of cultured C2C12 myoblasts are between 12-15 kPa (Truskey GA et al, 2002, Discher DE et al. 2004). Discher et al.2004 demonstrated that matrix stiffness of 12-15 kPa exhibits actual F actin assembly, moderate vinculin enriched focal adhesion in C2C12 cells, and myosin/actin striation in C2C12 myotube. Hence, according to the reports above, substrate elasticity has a significant impact on cell fate.

500 mutations in the LMNA gene have been discovered to be associated with the pathogenesis of 16 different diseases, primarily affecting muscle and adipose tissues (Capell,Collins B.C 2006, Bonne G, Muntoni F 2000, Raffaele Di Barletta, Toniolo D. 2000). A mouse lacking LMNA dies prematurely as a result of the severe muscular dystrophy phenotype ( Burke B et al. 1999). Lamin A is the prime component of a nucleus that is affected by tissue stiffness and matrix elasticity (Swift, J. Discher, D.E 2013; Harada, Discher D.E. 2014). In laminopathies the nucleus is severely damaged as a defining feature (Ho C,Lammerding J 2013). Lamin A was thought to function as a mechanosensor by interacting with the LINC complex, which aids in force transmission by changing gene expression patterns in response to changes in cellular environments (Fedorchak GR, Lammerding J. 2014). Lamin A null mouse embryonic fibroblasts were previously tested for nucleus elasticity and load bearing capacity using AFM indentation, micropipette aspiration, and microrheological experiments (Radmacher, M. 2007,Rowat, Aebi U. 2008). Loss of lamin A reduces the stiffness and fragility of the nucleus (Burke B et al. 1999, Lee RT et al. 2006), resulting in a mechanically weak nucleus. According to previous research, lamin A regulates the organisation of the cytoskeletal system. In LMNA −/− mice myocytes, overall cell shape is altered due to decreased connectivity with desmin filaments at the nuclear periphery (Fatkin D et al. 2004). Missense mutations in Ig fold lamin A that cause EDMD (R527P and L530P) result in 66% less F-actin binding (Wilson KL et al. 2010). The absence of lamin A causes significant disruptions in the assembly of vimentin, tubulin, and actin (Ramaekers FC et al.2004). In LMNA−/− MEFs, significant changes in cytoplasmic rheology are observed (Wirtz D et al. 2007). The LMNA mutant (N195K) cell model alters actin polymerization as well as the actual nucleo-cytoplasmic shuttling of a cytoskeletal protein regulator (Lammerding J et al. 2013). Previous research suggests that the interaction of the nucleoskeleton and the cytoskeleton is involved in the transmission of external mechanical cues to the nucleus. Actin filaments form a perinuclear actin cap in a pre-stressed state to keep the nucleus stable (Khatau, Wirtz D. 2009; Versaevel, M;Gabriele S. 2012, Mazumder, Shivashankar G.V 2008). Changes in the cytoskeleton affect the location of microtubule organising centres (MTOC) and the cell’s migration speed (Wirtz D et al. 2007). Central and apical stress fibres in endothelial cells balance the mechanistic relationship between cell and nucleus (Versaevel, M;Gabriele S. 2012).

Taking cues from the previous findings, we used a 12-15 kPa smooth surface and a 12-15 kPa engineered pattern surface to elucidate the effect on cell homeostasis, which is still poorly understood as a result of muscular dystrophy mutations. LA R453W and LA W514R mutations were chosen based on the severity of the phenotype (www.umd.be/LMNA).

According to the mutational database, 66 records have been reported for the occurance of R453W mutation in exon 7 of LMNA, associed with Emery Dreifuss Muscular Dystrophy (EDMD). Where as, for the W514R mutation, 3 records have been cited so far in exon 9 of LMNA gene, implicated in skeletal muscular dystrophy. Contractures are common in the early stages of muscular dystrophy, followed by muscle wasting and cardiac conduction defects (Emery AE, Dreifuss 1966, M Vytopil,D Toniolo 2003). To determine the molecular basis in the phenotypes of muscular dystrophy mutations, we used two approaches. First, we thoroughly examined cell and nuclear morphology, as well as the distribution of actin and focal adhesion complexes. Second, transcriptome analysis was used to confirm gene expression levels and regulators.

## Results

### Differential cell adhesion and shape on pattern surface

Substrate modulates cell adhesion and spreading (H Kenar et al.2006, Pepe-Mooney BJ et al. 2012). The contact area has a significant impact on cell confinement. We cultured the same number of cells (15000 cells/well) on a 12-15kPa smooth and pattern substrate with a gap of 10 μM between two patterns and a height of 400 nm. We discovered that fewer cells grew on the pattern substrate (**Figure 1 -a,b**). This explains the pattern substrate might have provided less contact area, which can regulate cell adhesion. However, cell alignment with respect to the pattern’s axis was not observed in some cases, which contradicted the findings of Brunette DM et al.1993, Iwata H et al.2009, and Borenstein JT et al.2009. These differences could be evolved due to the low height of the pattern substrate used in our study.

**Figure 1:**
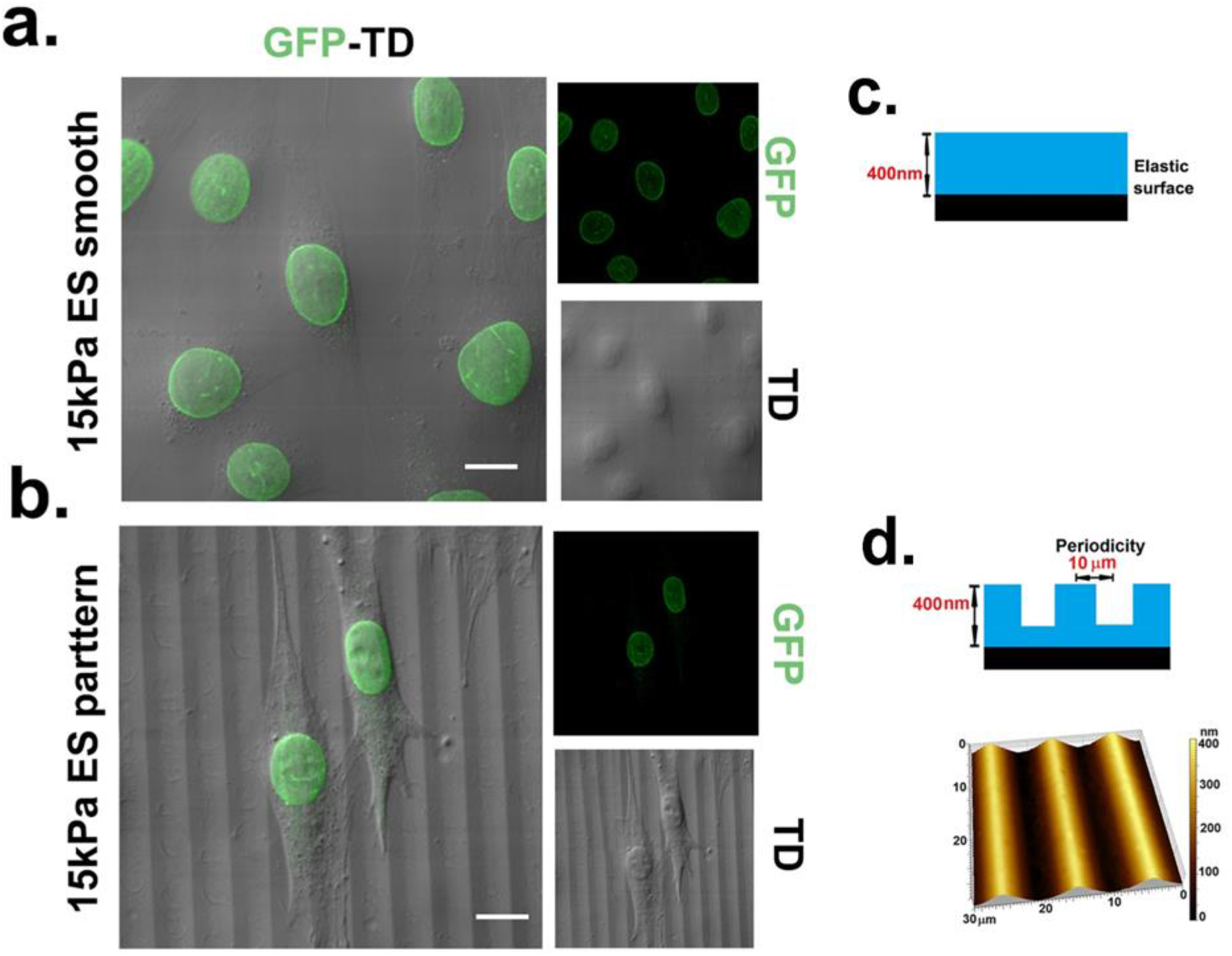
Cell attachment layout on distinct surfaces: (a),(b) Cell density on a specific field of ES smooth, ES pattern respectively, when started with same cell no. 15000 cells/ well and incubated for 24 hr.(d). AFM image of ES pattern surface.

AFM analysis was performed in accordance with the preceding findings (**Figure 2**). Cell shape parameters were calculated using the formulae depicted in method section. Table.1 contains information on cell morphology derived from AFM studies. On the pattern substrate, cells had a higher aspect ratio, eccentricity, and lower circularity. When compared to the wild type, mutants showed a lower area, perimeter, circularity, and a higher aspect ratio and eccentricity.

**Figure 2:**
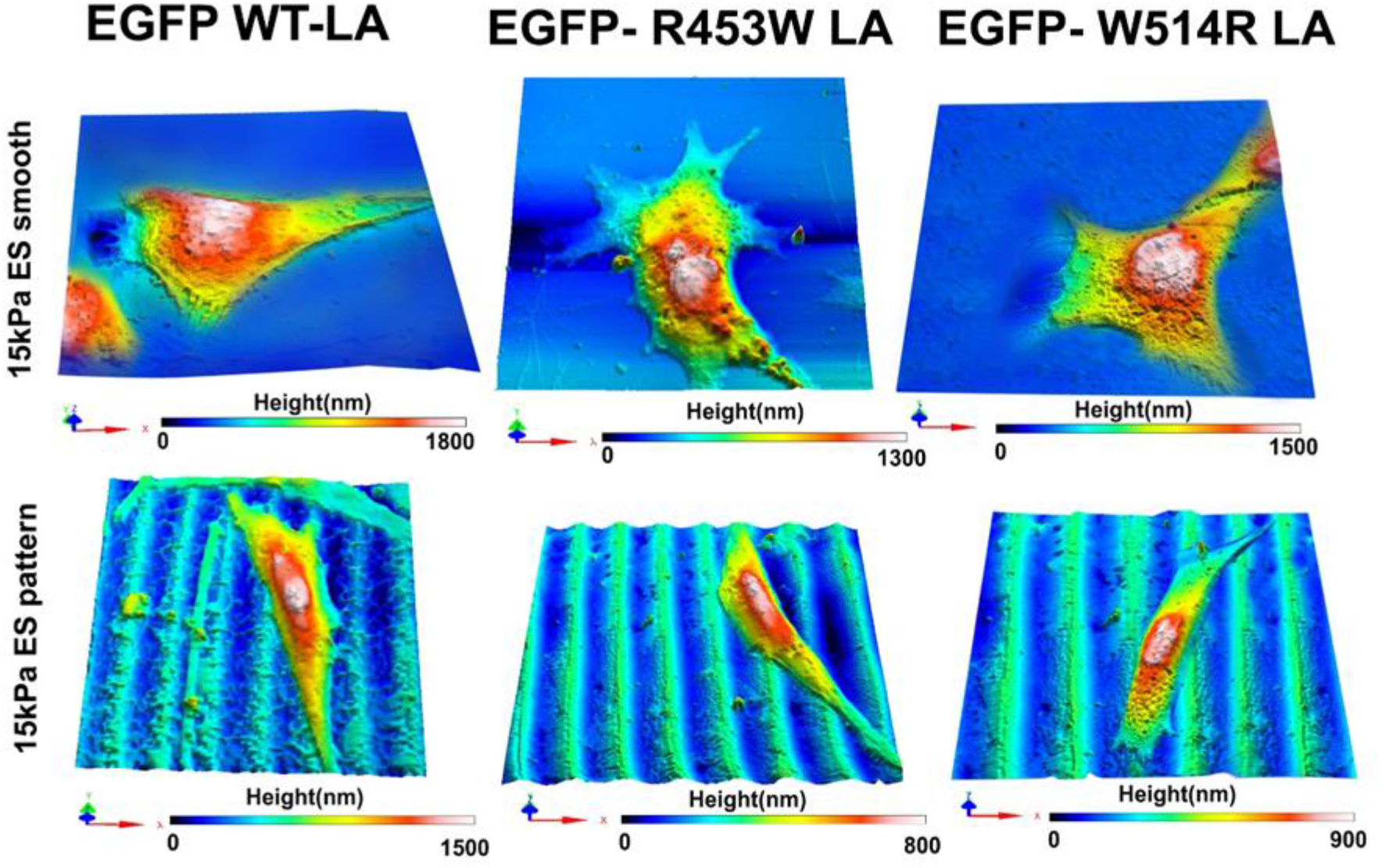
3D AFM images of EGFP WT LA, EGFP R453W LA, EGFP W514R LA. Corresponding histograms define maximum height of the cell.

**Table 1:** Calculations of cell morphometry parameters.

| Parameters | WT LA on 15 kPa ESS | R453W LA on 15 kPa ESS | W514R LA on 15 kPa ESS | WT LA on Pattern | R453W LA on pattern | W514R LA on pattern |
| --- | --- | --- | --- | --- | --- | --- |
| Aspect ratio | 3.17±.39 | 3.90±.48 | 4.38±.50 | 3.81±.47 | 5.39±.62 | 5.45±.67 |
| Eccentricity | 0.948±.11 | 0.966±.14 | 0.973±.10 | 0.965±.12 | 0.982±.13 | 0.982±.9 |
| Area | 6212.37±388 | 4900.88±306 | 4461.12±278 | 5805.66±362 | 4178.84±261 | 4107.18±256 |
| Perimeter | 385.79±52 | 382.6±47 | 381.24±45 | 368.68±46 | 357.76±44 | 359.5±43 |
| Circularity | 0.57±.07 | 0.48±.05 | 0.43±.02 | 0.49±.04 | 0.35±.03 | 0.35±.02 |

### Nucleus deformation as a result of pattern surface

The nucleus of the cell have a tendency to deform because of the microstructured surfaces (Davidson, Anselme 2009). We discovered slightly elongated nuclei for the introduction of pattern surface through immunofluorescence studies (**Figure 3-b**). In this context, recent studies by Weiss and Garber.1952, Khatau et al.2009, Versaevel et al. 2012 unveiled that nuclear morphology is consistent with cell morphology, as from their study they discovered elongated cells typically display an elongated nucleus morphology. In addition,AFM height map analysis revealed that the height of the cell on the pattern surface was reduced, lending credence to the elongation of nuclei on the pattern surface. On a 12-15 kPa smooth elastomeric surface, LA R453W cells induced a bleb-like structure with aggregates, whereas LA W514R cells presented lamin protein aggregation rather than a nuclear rim-like structure of wild type cells (**Figure 3- a,b**). When mutant lamin A expressed cells were grown on a pattern surface, we found more aggregations in the nuclei than mutant cells grown on a smooth surface.

**Figure 3:**
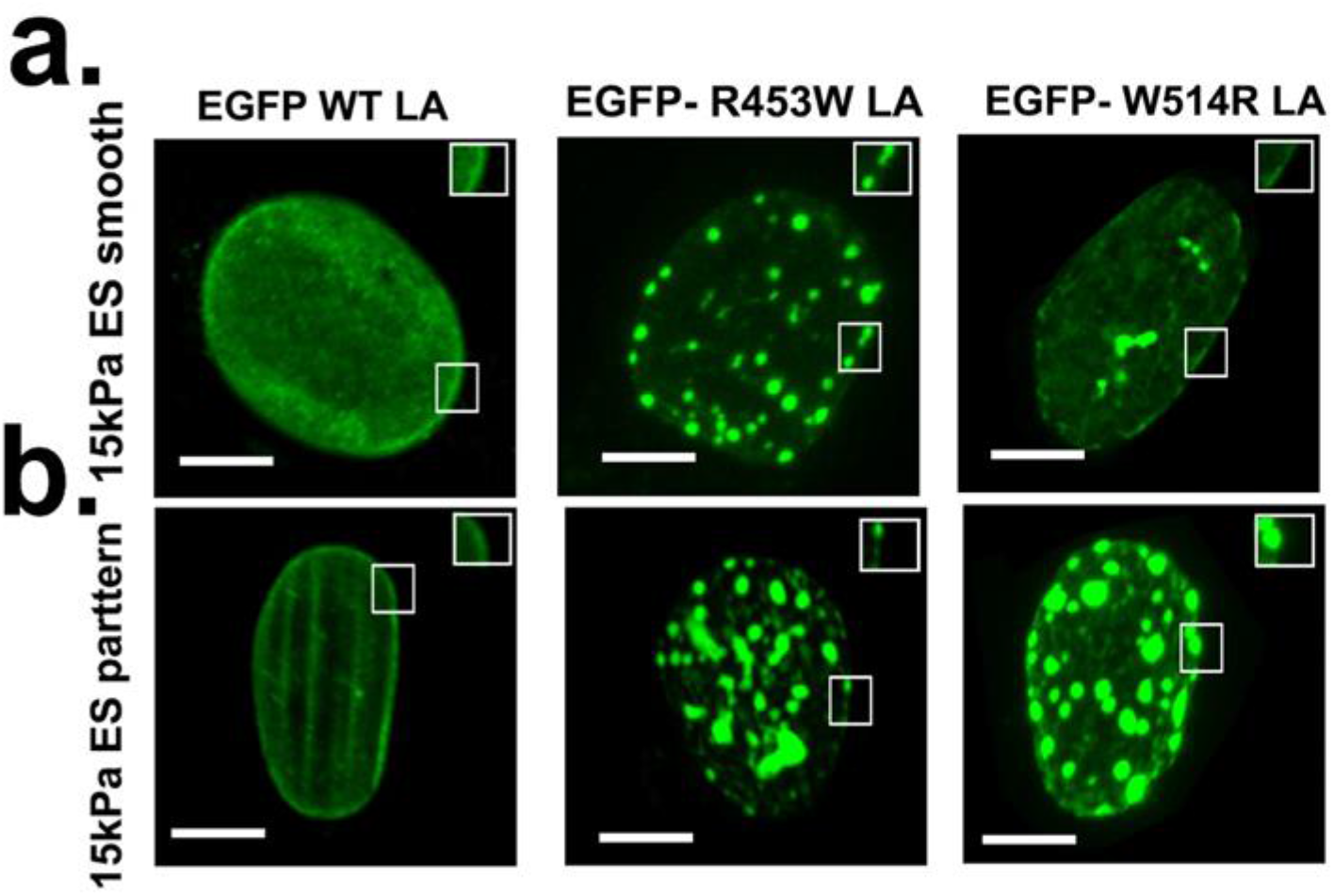
Confocal micrographs presenting distribution of lamin A in wild type and mutant cells on different surfaces. (a). nucleus morphology on smooth surface. (b). nucleus morphology on pattern surface. Mutant cells promote deformation of nucleus in maximum projection. Scale bar = 5μm.

### Distribution of actin and focal adhesion protein in response to pattern surface

The cytoskeleton detects and processes any extracellular force in the form of biochemical signals and mechanical forces that enter the nucleus for further downstream processing ( Schwartz MA et al. 2011).Surprisingly, the immunofluorescence analysis revealed the presence of actin monomer like structures in cells as a result of the mutations (**Figure 4-b**). This discovery was comparable to wound closure in Xenopus oocytes (Mandato & Berment,2001). When compared to the wild type, we found that actin monomers were more prevalent in LA R453W expressed cells than in LA W514R expressed cells. More actin monomer like structures were found on the pattern surface than on the smooth elastomeric surface. In both mutants, we found less dense actin networks than WT LA cells. Based on RNA sequencing analysis, we discovered a low level of ACTA1 and ACTC 1 genes, which confer reduced actin polymerization in mutant cells (**Figure 4-c,e**). Interestingly, high impediment in the expression of these genes were displayed in mutant cells on the pattern surface. Together, our analyses confirm that actin stress fiber formation is dependent on the matrix environment (Kim DH et al. 2012). These results are depicted by a cartoon in **Figure 4-d**.

**Figure 4:**
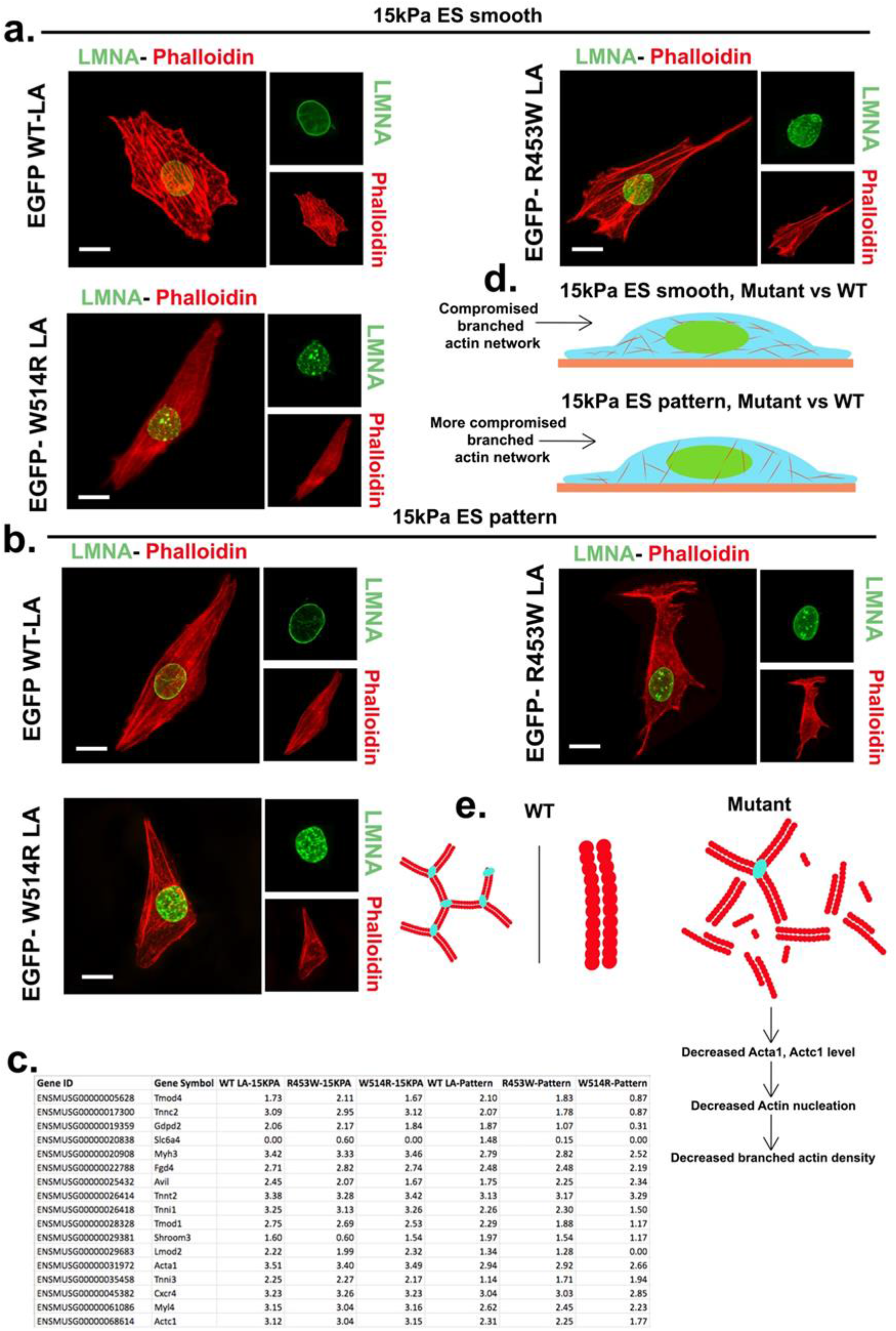
Lamin A mutations lead to differential status of actin on different surfaces. (a),(b) are Alexa Fluor 568 –phalloidin stained cells on smooth surface and pattern surface respectively. R453W LA, W514R LA respectively show less dense actin filamentous network with actin nodes. Actin nodes formation are higher in mutant cells on pattern surface. Scale bar = 10 μm.(c). cartoon depicts an illustration of comparative findings about actin filament networks among wild type and mutant. (d) Table of gene expression levels of GO term “Cytoskeleton”. (e). cartoon show regulatory factors influencing decreased branched actin density between wild type and mutant.

Actin changes could reflect the distribution of mechanosensitive proteins (Colombelli et al. 2009).In our study, we chose Vinculin, a major candidate of focal adhesion complex that exerts approximately 30% of the adhesion strength of cells when a 200 nN adhesive force is applied (Gallant et al. 2005). LA R453W and LA W514R had lower vinculin protein counts at the cell periphery than WT LA. We found fewer focal adhesion proteins, such as vinculin, at the adhesion plaque on the pattern surface (**Figure 5-a,b**). We discovered dramatic dysregulation of the following GO pathways from RNA sequencing analysis: extracllular space, extracellular region, cell junction, proteinaceous extracellular matrix, homophilic cell adhesion via plasma membrane adhesion molecules, extracellular matrix, collagen trimer, collagen degradation, basement membrane, assembly of collagen fibrils and other multimer structures, extracellular matrix degradation, collagen binding. In mutant cells on pattern substrate, we found more downregulation (as well as when compared to the upregulated genes) of the GO term: extracellular (**Figure 5-c,d**). S100b, Mrgprd, and Pcdhb6 genes were found to be downregulated. Furthermore, qRT PCR results revealed a low gene expression level of Focal Adhesion Kinase (FAK) and two important isoforms of Integrin (Integrin 6 and Integrin 1) for the pattern surface in comparison to a smooth substrate (**Figure 6**).

**Figure 5:**
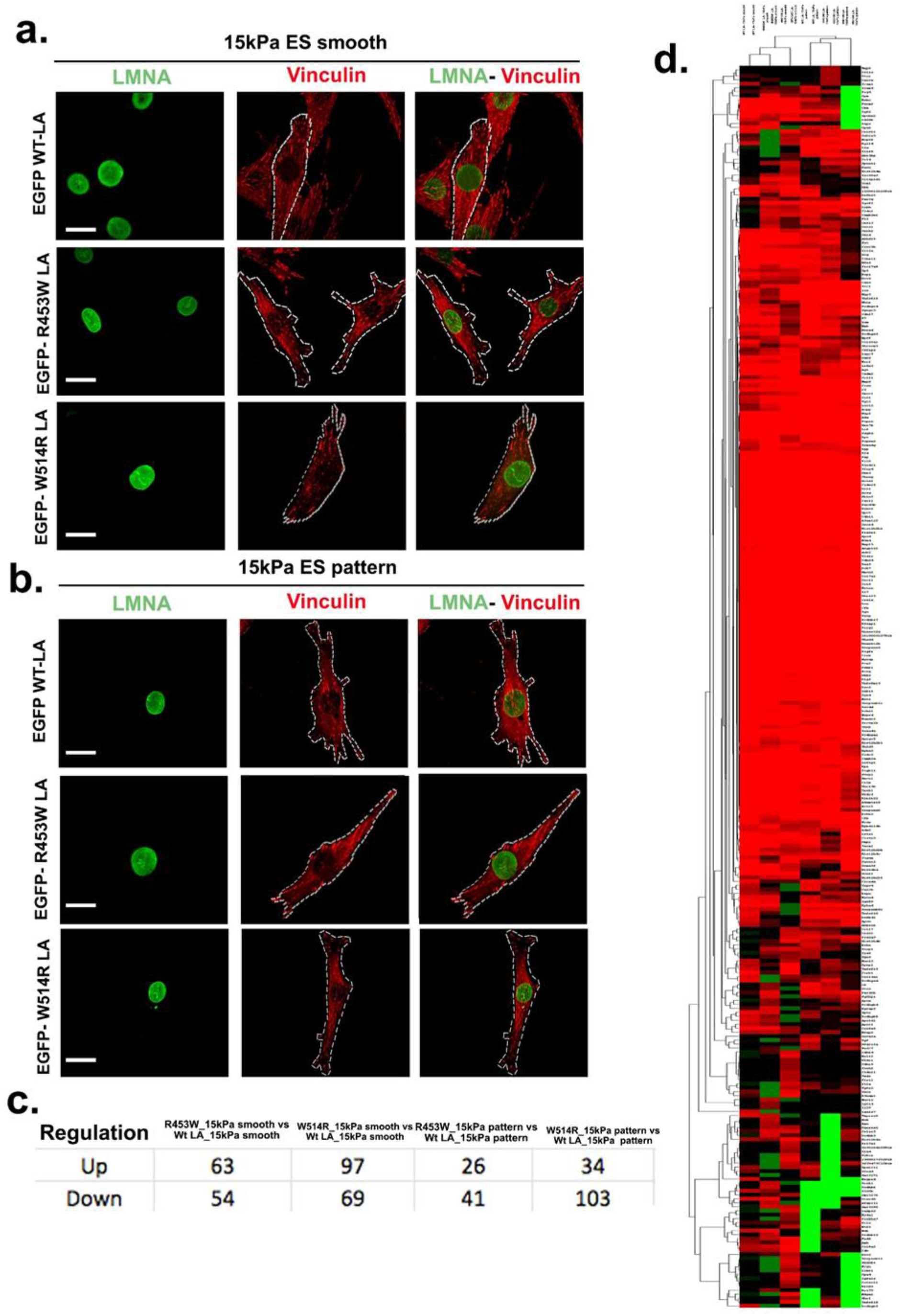
Anomaly in the vinculin recruitment and other components of extracellular signaling in the periphery of lamin A mutant cells. (a),(b) Vinculin is stained in red (Alexa Fluor 568) in stably transfected cells on smooth and pattern surface respectively. Boxed regions indicate more vinculin accumulation inside the nucleus of mutant cells irrespective of surface. (c). upregulation and downregulation gene numbers in GO term “extracellular”. (d). heatmap of genes present in GO term “extracellular”.

**Figure 6:**
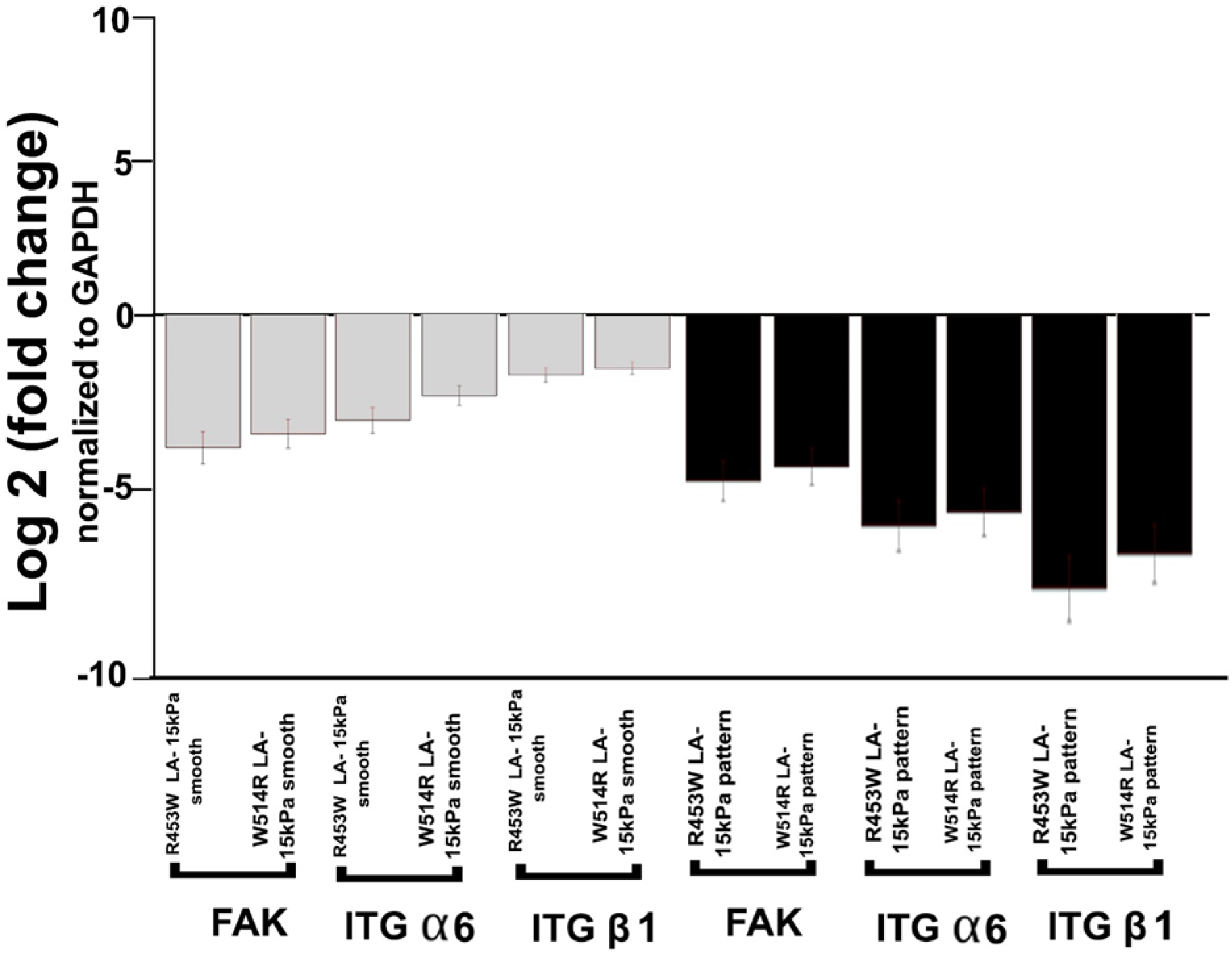
Expression patterns of selected genes in lamin A mutant cells determined by qRT-PCR. qRT PCR analysis for FAK, ITG α6,ITG β1in response to the 2 experimental conditions of substrate. The gene expression was normalized by the gene expression of GAPDH and statistical analysis was also performed.Log 2 value of fold changes were used for bar graph plotting.

### Alteration of LINC complex on pattern surface

The LINC complex is a critical component in the mechanotransduction cascade (Hodziz D et al. 2006). SUN1 and SUN2 proteins as well as the Nesprin protein family are found in the LINC complex. The inner nuclear membrane proteins SUN1 and SUN2 can interact directly with the lamin A protein. Nesprin proteins bind to SUN1/2 via their C terminal KASH domain (Klarsicht,Anc—1,Syne homology) and interact with Actin binding domain via their N terminal domain. Smith JD et al.2019 demonstrated that SYNE2 overexpression reduces nuclear stiffness along with nucleus enlargement and cytoskeleton displacement by decreasing the level of Nesprin 2 giant isoform. During meiosis, the SUN1–ZYG12 LINC complex attaches chromosomes to the nuclear envelope and aids in essential pairing and crossover of homologous chromosomes, resulting in the formation of a Holliday junction (Fridkin A et al. 2007, Machacek T et al. 2009, Carlton PM, et al. 2009). SUN1 (Han M et al. 2007), SUN2 (Benavente R et al. 2014), and KASH5 (Stewart CL et al. 2013, Ishiguro K et al. 2012) tether telomeres on the lamina to enable the above roles in mammalian meiosis.

RNA sequencing analysis revealed that mutant cells on the pattern surface had down regulation of SUN1 and SUN2 genes expression along with slightly differential expression of SYNE2 gene in the mutant cells(**Figure 7**). Interestingly,Chen et al. (2012) observed SUN1 overexpression in lmna−/− mice.

**Figure 7:**
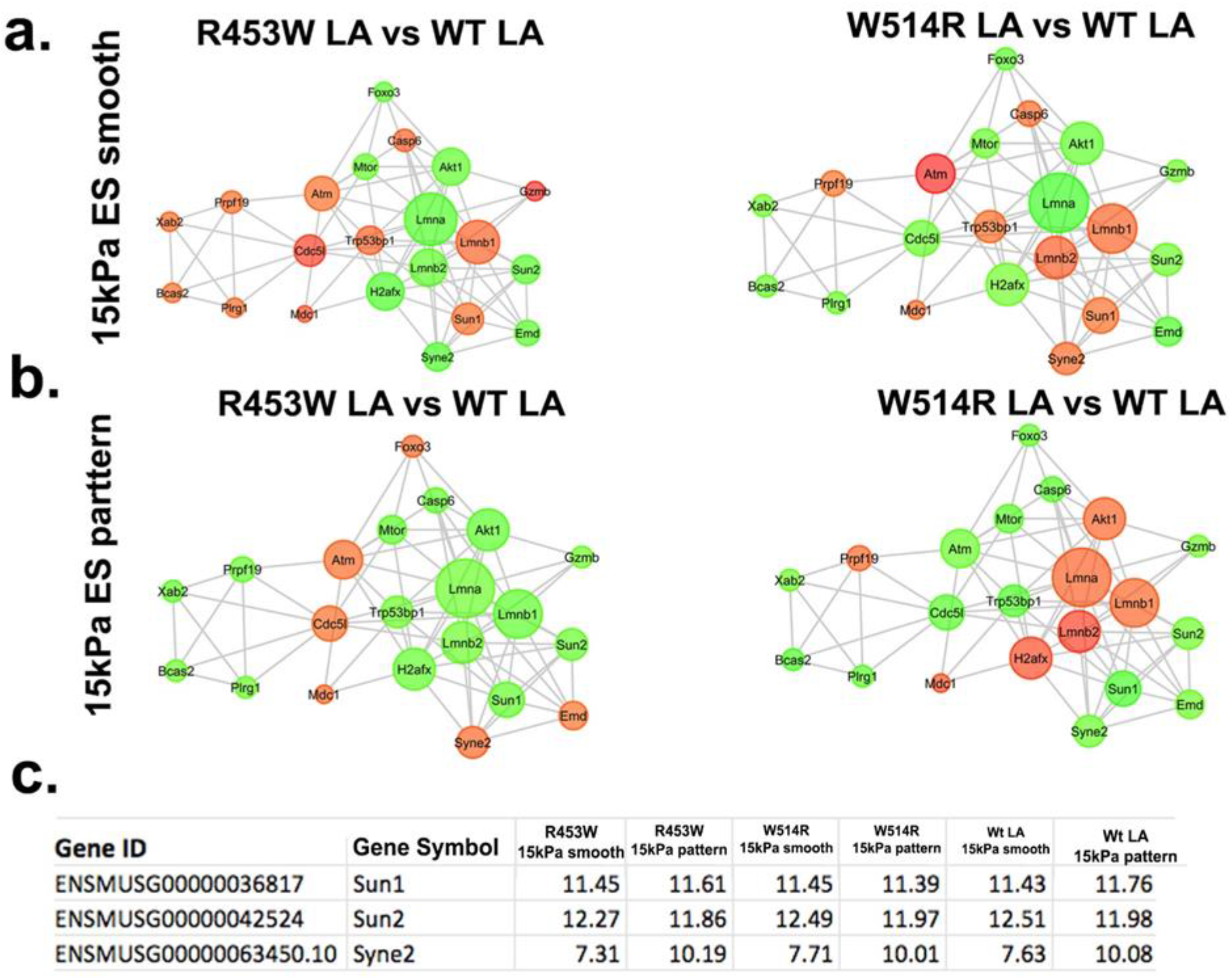
The expression characteristics of genes of LINC complex in wild type and lamin A mutated cells. (a),(b) Lamin A interactome analysis for R453W LA vs WT LA, W514R LA vs WT LA when grown on ES smooth and ES pattern. (c). Expression levels of SUN1, SUN2, Syne2 in RNA sequencing.

### Functional categorization of static stimuli treated WT-LA and mutant-LA DEGs

Comparative analysis of the transcriptomes of WT and the mutants of Lamin A associated muscular dystrophy indicates impairment in the expression of multiple gens and pathways under different static stimuli. The data related to GO enrichment and KEGG pathway analysis will be uploaded shortly.

Decoupling of nuclear-cytoskeletal machinery acts as an observatory for the topology of chromosome (Fedorchak et al.2014, Le et al. 2016, Maharana et al.2016, Shachar and Misteli et al. 2017) which may affect the global gene expression to sustain mechanical loading of pattern surface. Results of Condition tree, PCA (Principle Component Analysis) and Correlation Matrix using Cluster 3.0 tool showed significant reproducibility among the samples within the same conditions (12-15 kPa smooth surface and 12-15 kPa pattern surface) and distance between the groups. Further, the resolved transcriptome showed 26466 genes were detected in either of the groups. Differentially expressed transcripts were identified using Cuff Diff pipeline (REF) with a Fold Change cutoff of >=2.0 and a student t-test p Value of <=0.05. This resulted in identification of reversal of gene regulation in LA W514R mutant when cultured in smooth vs pattern, where in smooth surface there was more of up regulation and in pattern there was more of down regulation. This is not true in case of R453 mutant samples. Differential gene expressions were visualized using volcano plot and unsupervised hierarchical clustering. Genome wide distribution of DEGs of mutants showed very poor enrichment in chromosome 10,12,16 and Y. Venn Diagram based analysis of DEGs between mutants in 15 kPa Pattern and 15KPa smooth showed inverse regulation of 31 genes between LA R453W on 15kPa smooth and LA W514R on 15kPa smooth. Primary analysis on alteration in gene expression profile due to mutations in lamin A were presented in **Supplementary Figure 1**. GO and pathways as analyzed by DAVID tool with a FDR score of <=0.05 was considered for further downstream analysis. This resulted in 92 unique GO categories and Pathways from multiple databases that harbor the differentially expressed genes. Phenotype vs Genotype analysis was done to identify statistically significant and enriched GO and Pathways, which resulted in identification of 12 non-redundant GO and Pathways. 12 GO and pathways that are involved, are: ion transport, DNA damage & repair, Muscle gene regulation, cytoskeleton, chromatin organization, senescence,signal transduction pathway,immune response,epigenetics & transcription regulation, meiotic recombination and synapsis,DNA binding, extracellular regions etc (**Supplementary Figure 2**). As per gene regulatory network analysis from 12 GO terms, LA W514R possessed a gross molecular changes than LA R453W mutant cells due to the variation in surface property ( **Figure 8**). Expression of LA R453W, LA W514R in cells on pattern surface promoted highly compacted state on chromatin by means of upregulating the epigenetic genes. Consequently, direct readout of high chromatin condensation demonstrated repression of muscle genes transcription. Additionally, heatmap for genes involved in Signaling cascades were also made to present the expression of key gene pools (**Supplementary Figure 3**).

**Figure 8:**
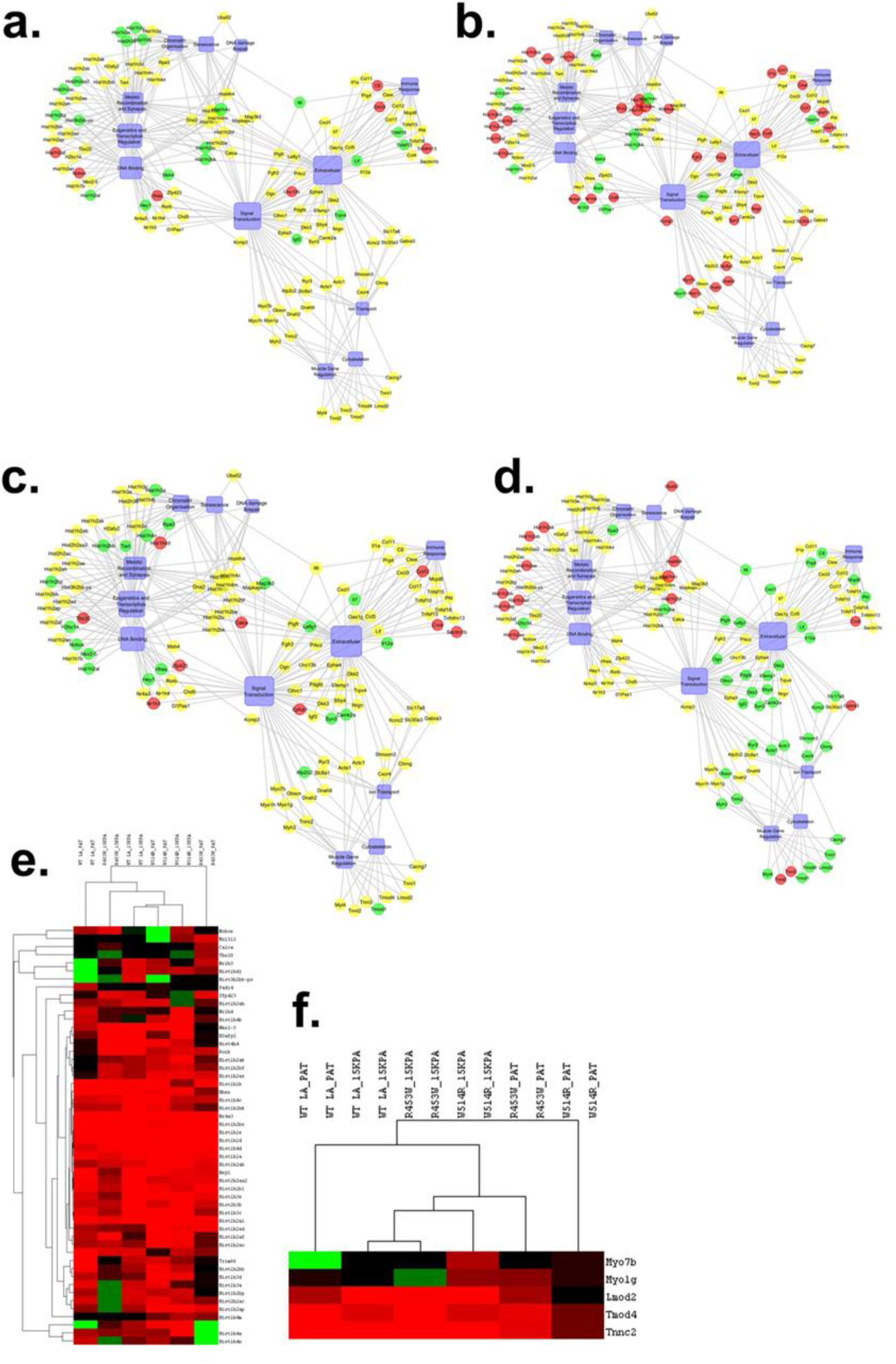
Network analysis unveils deregulated regulatory circuits and overlap among the pathways in lamin A mutant cells upon the introduction of distinct surfaces. (a), (b), (c), (d) Gene regulatory networks of R453W LA (smooth surface), W514R LA 9 smooth surface), R453W LA (pattern surface), W514R LA (pattern surface). Red sphere indicates upregulation, green sphere indicates downregulation. (e),(f) are heatmaps of GO term “epigenetics & transcription regulation” and “muscle gene regulation”.

## Discussion

The nucleus, which is typically spherical or ellipsoidal in shape, is the stiffest organelle of the cell, being 2-10 times harder than any other cytoskeletal elements (Meister JJ et al.2002, Burgkart R et al. 2000). However, physical or environmental changes, forces, and strain can cause dramatic morphological changes (Aebi U et al. 1986, Dahl K et al. 2008,Roca -Cusachs et al. 2008,Dauer and Worman 2009, Hampoelz et al. 2011,Gerlitz et al. 2013). Nuclear lamina, which is made up of lamins, is crucial in maintaining the structural integrity of the nucleus (Discher DE et al. 2004,2005). Using lmna-deficient or mutant cells as model systems, Lammerding et al. 2004,2005, Hale et al. 2008, Khatau et al. 2012 asserted that matrix stiffness, topography, and dimensionality can serve as observatories to better understand nuclear mechanics and cell mechanotransduction. In this study, we attempted to mimic muscle-like physiological environments in order to gain a detailed understanding of the plasticity of the cytoskeleton, nucleus, and focal adhesion complexes in LMNA mutated myoblasts. Our current findings show that this is the first report that combines structural and gene expression hypotheses to propose a potential mechanical code for the pathogenesis of lamin A associated muscular dystrophy under physiological conditions. These findings are a clear example of lamin A’s expanded role that extends far beyond the nucleus. We also compared the equilibrium states of cells on the flat/smooth and pattern surfaces.

The purpose of this study was to determine the role of lamin A in the maintenance of coordination among the components of cellular mechanotransduction, when subjected to various engineered physiological conditions. We observed deformed nuclei with aggregates as a result of mutation because lamin A is a dominant regulator of nuclear shape. It is worth noting that the localization of nuclear aggregates on the pattern substrate is more in the nucleoplasm than in the nuclear periphery. As proposed by Taimen P et al. 2016, a possible explanation would be defective assembly machinery in lamin proteins as a result of mutation. They reported that an increase in solubility caused by mutation, as well as a disruption in the assembly machinery, could be the causes of nucleoplasmic aggregate formation. In addition to their observations, we can assume that the pattern substrate exerted more mechanical cue than the smooth substrate, resulting in more nuclear aggregate deflection toward nucleoplasm. Furthermore, the introduction of a pattern surface causes nuclei to elongate, indicating softening of the nuclear lamina due to muations. We also discovered that the mutant cells on the pattern surface had lower heights. Hock J et al. 1998 proposed that when a cell attaches to a micropillar, it pulls the nucleus downwards. In agreement with this previous report, we can say that cells are encountering low contact area due to the distinct surface topography of the pattern surface. For this, they require more forces for adhesion on this particular surface, which could explain the decrease in mutant cell heights on the pattern surface. Consequently, low cell-matrix contact area has a strong influence on the degree of cell spreading, and as a result, we observed a reduction in the area and perimeter of the cell on the pattern substrate. The circularity value should be 1.0 for a perfect circle, and if it approaches 0.0, it indicates a conversion to an ellipse. On the pattern substrate, mutant cells had a high aspect ratio and eccentricity, as well as a lower value of circularity. Previous research has shown that the attachment and spreading of a cell on a substrate is dependent on both inward and outward stresses experienced by the cell, and that these stresses ultimately define the shape and volume deformation of the cell (Kumar S.et al.2006, Dubin-Thaler BJ et al.2008, Griffin, M. A. et al. 2004, Wang N. et al. 2002, Ingber DE et al.1998,2002). As a result, our calculated parameters of cells (wild type cells and mutant cells) on both substrates indicate that low contact area may have induced mutant cell elongation due to disturbances in whole cell elasticity. Furthermore, these cell parameters establish a linear relationship between the shape of the cell and the shape of the nucleus. In this regard, we looked into other biochemical fractions of the cell besides lamin A. Previous research suggested that improperly assembled actin networks serve as a testament to actin’s regulatory property in the deformation of the nulceus due to the application of stress (Maninova M, Vomastek T. 2014, Hyungsuk Lee, Kevin Kit Parker 2015). Wirtz D et al. 2005 discovered that actin fibre is solely responsible for the nucleus’s disk-like shape and inability to move out of the plane of focus during nuclear rotation. Similarly, we discovered impaired actin polymerization as well as a lack of stress fibres in mutant cells grown on pattern substrate. The absence of vinculin recruitment at the cell periphery in the pattern surface indicates a breakdown in the coupling between F actin and vinculin in channelling extracellular stimuli or force (Balaban et al. 2001; Ji et al. 2008). Our result showed reduced FAK and isoforms of Integrin gene expression levels for the mutant cells specifically on pattern surface, which further suggest reduced cell strength for extracellular matrix and thus, resulting in fewer mutant cells growing on pattern surface. In addition to this finding, we discovered a difference in gene expression levels of SUN1, SUN2, and SYNE2, indicating a defective LINC complex. Khatau SB et al. (2008) discovered that LINC complex impairment in a mouse model of laminopathies resulted in a weak cytoskeleton and adhesion features. Our findings show a complete transformation of the mechanotransduction cascade in mutant cells.Distinct surface topography of pattern surface indeed modulates the cell fate via poor mechanical couplings within participant molecules as more disorganization of mechanotransduction cascade were seen in mutant cells on pattern substrate. Furthermore, the disparity in gene expression levels found from RNA sequencing and qRT PCR results are consistent with previous reports demonstrating that decoupling of nuclear-cytoskeletal machinery acts as a regulator for chromosome topology (Fedorchak et al.2014, Le et al. 2016, Maharana et al.2016, Shachar and Misteli et al. 2017). As a result, our findings show that Lamin A has a significant impact on the remodelling of the cytoskeleton system and the maintenance of nuclear homeostasis in order to sustain mechanical loading on the pattern surface.

Defects in the mechanotransduction cascade, as described by Kennedy BK et al. 2006 and Lattanzi G et al. 2005, impair the ability of lmna−/− myoblasts to differentiate. Buendia B et al.2003 demonstrated that EDMD mutant expression impairs muscle differentiation process in C2C12 myoblasts. The process of muscle differentiation is beyond the scope of our research and must be recapitulated in order to combat fatal phenotypes in muscular dystrophy. Based on our findings, we can extrapolate that this cumulative plight in mutant cells could spearhead the inhibition of muscle differentiation process, as it would prevent necessary cell fusions. In this connection, in mutant cells we observed more downregulation of muscle specific genes due to pattern surface which could be regulated by tight packing of chromatin with the upregulation of histone genes.As a result, the onset of muscular dystrophy phenotypes would occur.

## Methods

### Site directed mutagenesis

All the mutations were generated using site directed mutagenesis methods in EGFP-LA plasmid and the details of the primers are reported in Bera et al.2014, Dutta et al,2018. The mutations were confirmed by the sanger sequencing method (Sanger,F, and Coulson, A.R. et al.1975).

### Stable cell line preparation

Mouse myoblast cell line C2C12 was first transfected with Lipofectamine 2000.Medium was replaced on the next day with G418 containing medium. G418 was titrated first and 850 μg of selection antibiotic G418 has been standardized to select stable clones. Medium of the dish was changed everyday for 3-4 weeks for stable cell line preparation.

### Cell culture

18×18mm coverslips with 12-15 kPa smooth elastic surface and 12-15 kPa patterned surface (between two patterns which has height of 400 nm and gap of 10 μM)[provided by Dr. Rabibrata Mukherjee, Department of Chemical Engineering, IIT KGP) were coated with 10 μg/ml fibronectin. EGFP-WT LA, EGFP-R453W LA, EGFP-W514R LA stably transfected C2C12 cells were seeded on coverslips and maintained in a humidified incubator at 37° C in 5% CO_2_.

### Cell fixation and microscopy

Mouse myoblast C2C12 cells either grown on 12-15 kPa smooth or patterned surface were fixed using 4% PFA for 10 min, followed by PBS wash. Cells were permeabilized by 0.2% Triton X-100 solution in PBS stained with Phalloidin Alexa Fluor 568(0.35 μM, Molecular Probe, Life Technologies), Anti-vinculin (V9131,Sigma) antibody of 1:500 dilution for overnight at 4° C temperature. Then cells were incubated with secondary antibody, Alexa 594 conjugate Invitrogen (A11005) for 2 hour at RT. Finally,after washed with washing buffer and PBS for 3 times for 3 minute coverslips were mounted on Vectashield (Vector laboratories) for imaging. Confocal images were captured by Galvano mode NIKON A1 RMP detector and Plan Apochromat VC 100x oil DIC N2 / 1.40 135 NA /1.515 RI objective with an additional 4x digital zoom in a NIKON Inverted Research Eclipse TiE Laser Scanning Confocal/NSIM microscope. A multi line Argon -Krypton laser (λ_ex_-457/488/561 nm) at 3% was used for green channel and a solid state laser (λ_ex_-561 nm) at 5% was used for red channel. For Z stacks a step size of 0.25 μm was maintained. 45 cells per sample were checked for the study.

### AFM Sample Preparation and Imaging

For AFM imaging of 4% paraformaldehyde fixed cells, contact mode AFM was performed using a Pico plus 5500 AFM (Agilent Technologies USA) with a piezoscanner maximum range of 100um. Micro fabricated silicon cantilevers of 450um in length with a nominal spring force constant of 0.02-0.77N/m were used from Nano sensors,USA. The cantilever resonance frequency was 6 - 21 kHz. The images (256 by 256 pixels) were captured with a scan size of between 0.5 and 5 um at the scan speed rate of 2 lines/second through Pico view 1.12 software. Images were processed by flatten using Pico scan software (Molecular Imaging Corporation, USA). All the images were presented in this report derived from the original data. Major axis, minor axis of the cells were measured manually using Pico scan 5 software. 10 cells per sample were estimated for the AFM experiment. Following formulas were used to calculate the cell morphology:

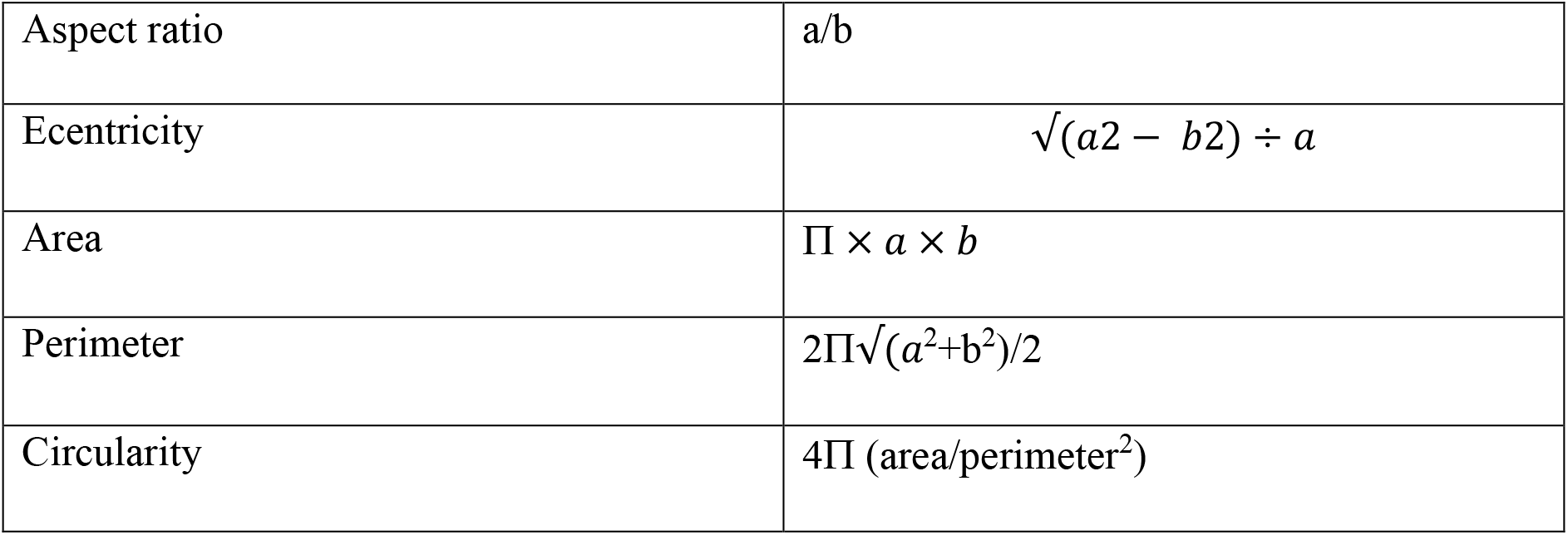

|  |  |
| --- | --- |
| Aspect ratio | $a/b$ |
| Eccentricity | $\sqrt{(a^2 - b^2)} \div a$ |
| Area | $\pi \times a \times b$ |
| Perimeter | $2\pi\sqrt{(a^2+b^2)}/2$ |
| Circularity | $4\pi (\text{area}/\text{perimeter}^2)$ |

### RNA isolation and sequencing

Total RNA was isolated by using spin column based RNA isolation kit (Qiagen). RNA quantity was determined by measuring absorbance at 260 nm using a NanoDrop UV-VIS spectrophotometer. 50 μg of total RNA was used as starting material for the preparation of whole transcriptome library. The yield and size distribution of the amplified DNA was assessed by running Bio analyser. After that, Ion Sphere particles (ISP) were prepared by Ion PI Hi-Q OT2 200 kit and ran on Ion PGM^™^ system.

### RNA sequencing data analysis

Raw Data obtained in FASTq format for the samples with a read length of 100bp, on an average provided 7.5 million HQ reads per sample upon quality control analysis using NGSQC Tool Kit. Further, the HQ passed reads were subjected to splice var alignment against mm10 Build GRCh38 using TopHat aligner, which resulted on an average of 85% of the HQ reads aligning to the reference genome. Analysis of degree of differential gene expression among the samples were done by plotting Condition tree, PCA (Principle Component Analysis) and Correlation Matrix using Cluster 3.0 tool. Further, the resolved transcriptome showed 26466 genes were detected in either of the groups. PartekFlow^™^ (Partek Inc,MO,USA) software was used to identify differentially expressed genes (DEG s) between wild type lamin A and mutant lamin A. Statistically significant differentially expressed transcripts were subjected to GO and Pathway enrichment using DAVID tool. Only those GO and pathways with a FDR score of <=0.05 was considered for further downstream analysis.

### qRT PCR

Details on this method was described in chapter 2. Briefly, total RNA extracted from the cells using Qiagen RNeasy mini kit. After that, cDNA was generated by Thermo Scientific RevertAid First Strand cDNA synthesis kit from 2.5μg of RNA. Using 100ng cDNA and two-step SYBR Green I RT-PCR master mix, qRT PCR was performed to detect SUN1, SUN2, FAK, Integrin α 6, Integrin β 1 by Agilent 2100 Bioanalyzer. All the experiments were performed in triplicates. The mean and standard deviation of the triplicates were taken and relative expression was calculated by the ddCT method. Post that, bar graph was plotted using Origin (OriginLab, Northampton, MA). We used GAPDH as internal control. Primers sets for the genes of interest are following:

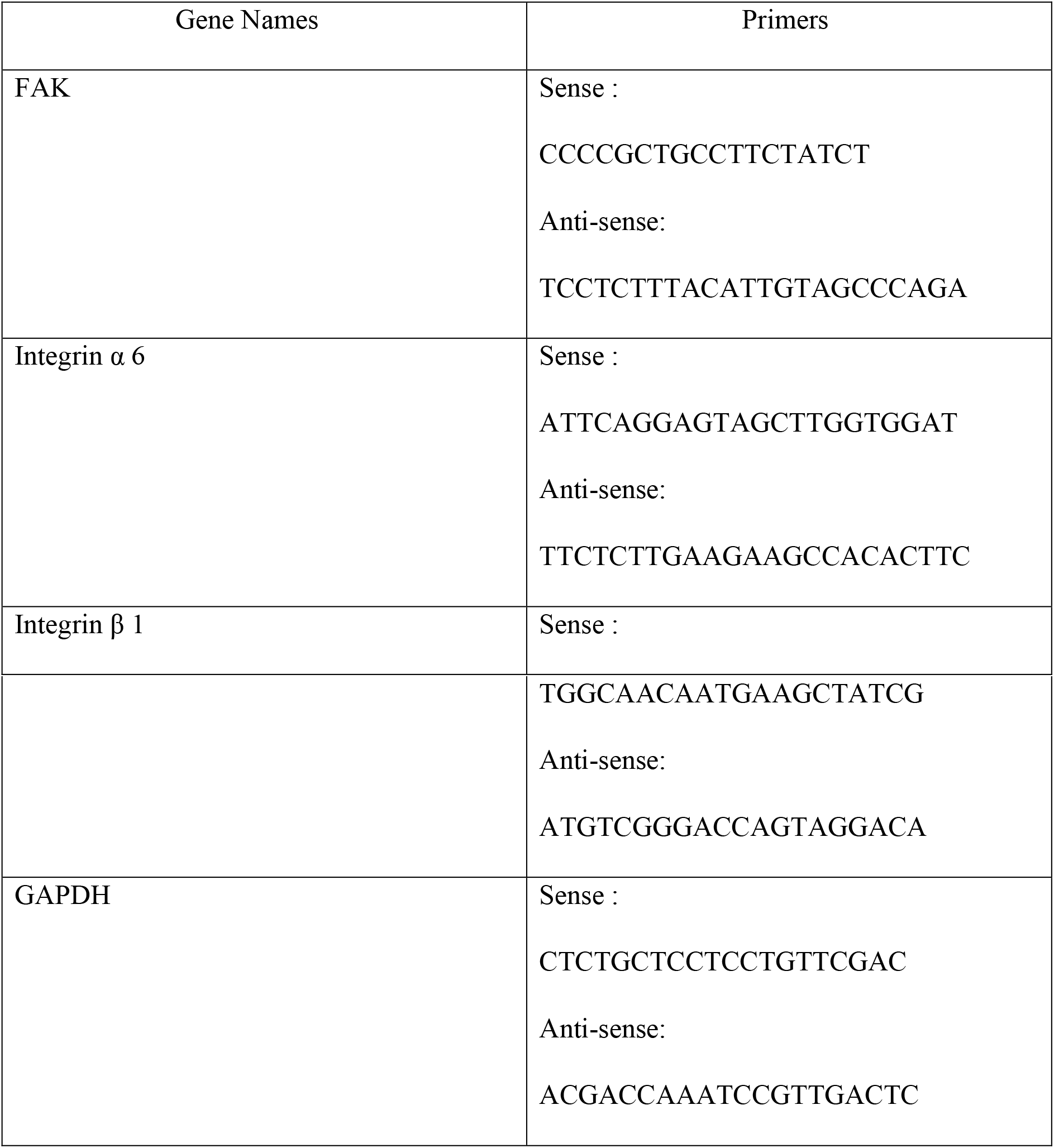

## Acknowledgments

The vectors pET-LA and pEGFP-LA were generous gifts from Dr. Robert D Goldman, Feinberg School of Medicine, Northwestern University,Chicago. We also thank Dr. Rabibrata Mukherjee (IIT KGP) for sharing of coverslips with different surface chemistry. For RNA sequencing authors acknowledge Mr. Saikat Mukherjee (Crystallography and Molecular Biology Divison, Saha Institute of Nuclear Physics), Ashish K.George (Thermo Scientific). Part of this research was funded by SERB-DST and BARD project,DAE.

## Author Contributions

S.D. conceived and designed the experiments. S.D. performed all the immunofluorescence and biochemical experiments used in this study. S.D., M.V. analysed the RNA sequencing data.S.D. and T.M. contributed in recording AFM and analysis. S.D. and M.V. make the draft of the manuscript. All authors have read and agreed to the published version of the manuscript.

## Supplementary Figures

**Figure Supplementary 1:**
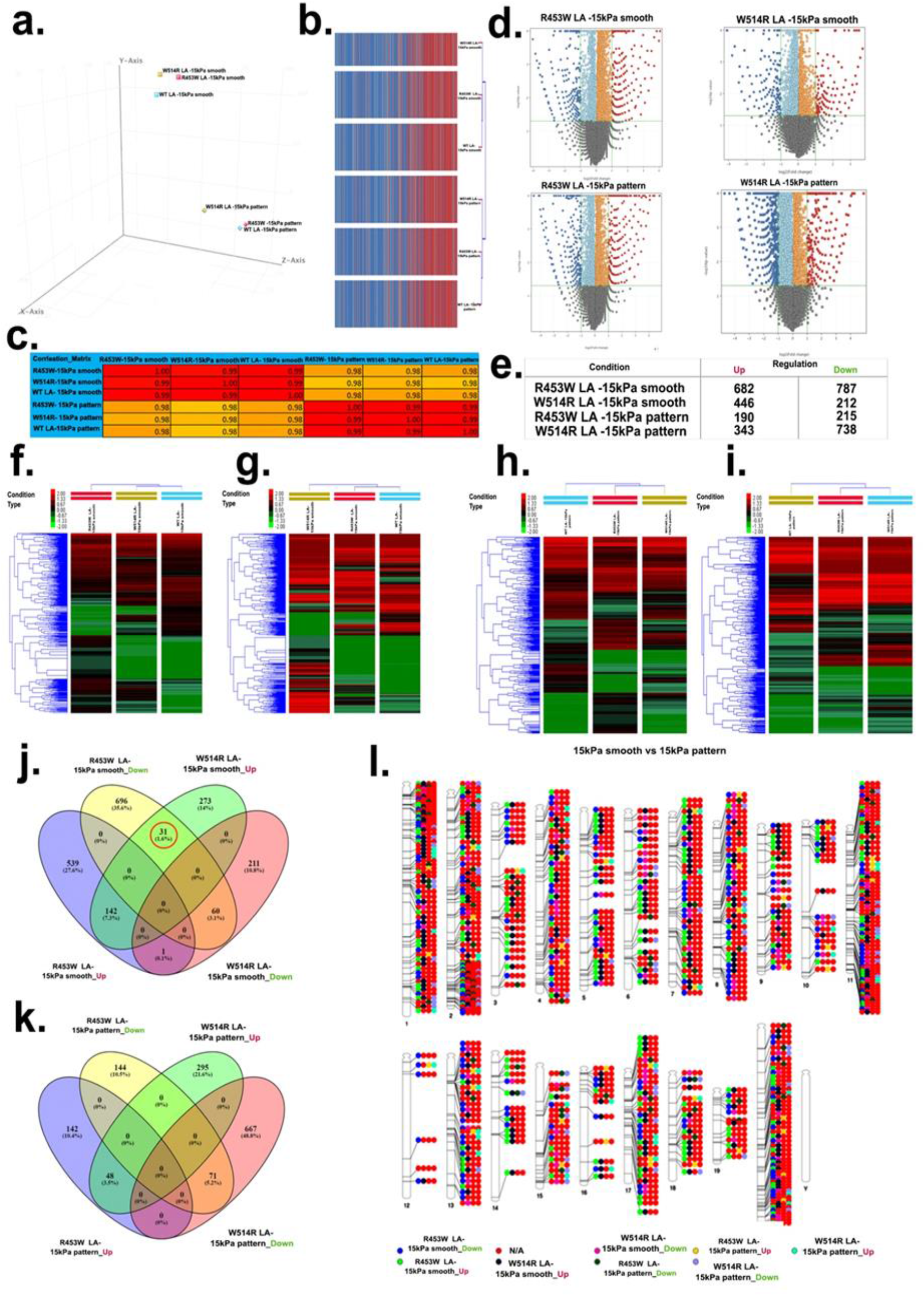
Qualitative and Quantitative Assessment of Deep Sequenced Transcriptome. (a), (b),(c) denote reproducibility among the samples. (a) PCA analysis, (b) Condition tree, (c) Correlation plot. (d) Volcano plots of each distinct mutant after the substitution from wild type on 2 surface conditions. (e) Table of regulated and down regulated genes of each samples.(f),(g),(h),(i) Unsupervised hierarchical clustering to figure out differentially expressed transcripts by Cuff Diff pipeline (REF). Fold Change cutoff of >=2.0 and a student t-test p Value of <=0.05 were set. Red color indicates upregulation, Black color indicates no regulation, Green color indicates downregulation. (j) and (k) Venn diagrams of R453W LA _ smooth surface vs W514R LA _ smooth surface and R453W LA _pattern surface vs W514R LA_ pattern surface illustrating the overlap in DEGs. The numbers of in each circle present the total number of different genes in each comparison group and the number in the overlapping areas is the number of shared gens between two comparison groups. (l)Genome wide distribution of DEGs of mutants via ideogram.

**Figure Supplementary 2:**
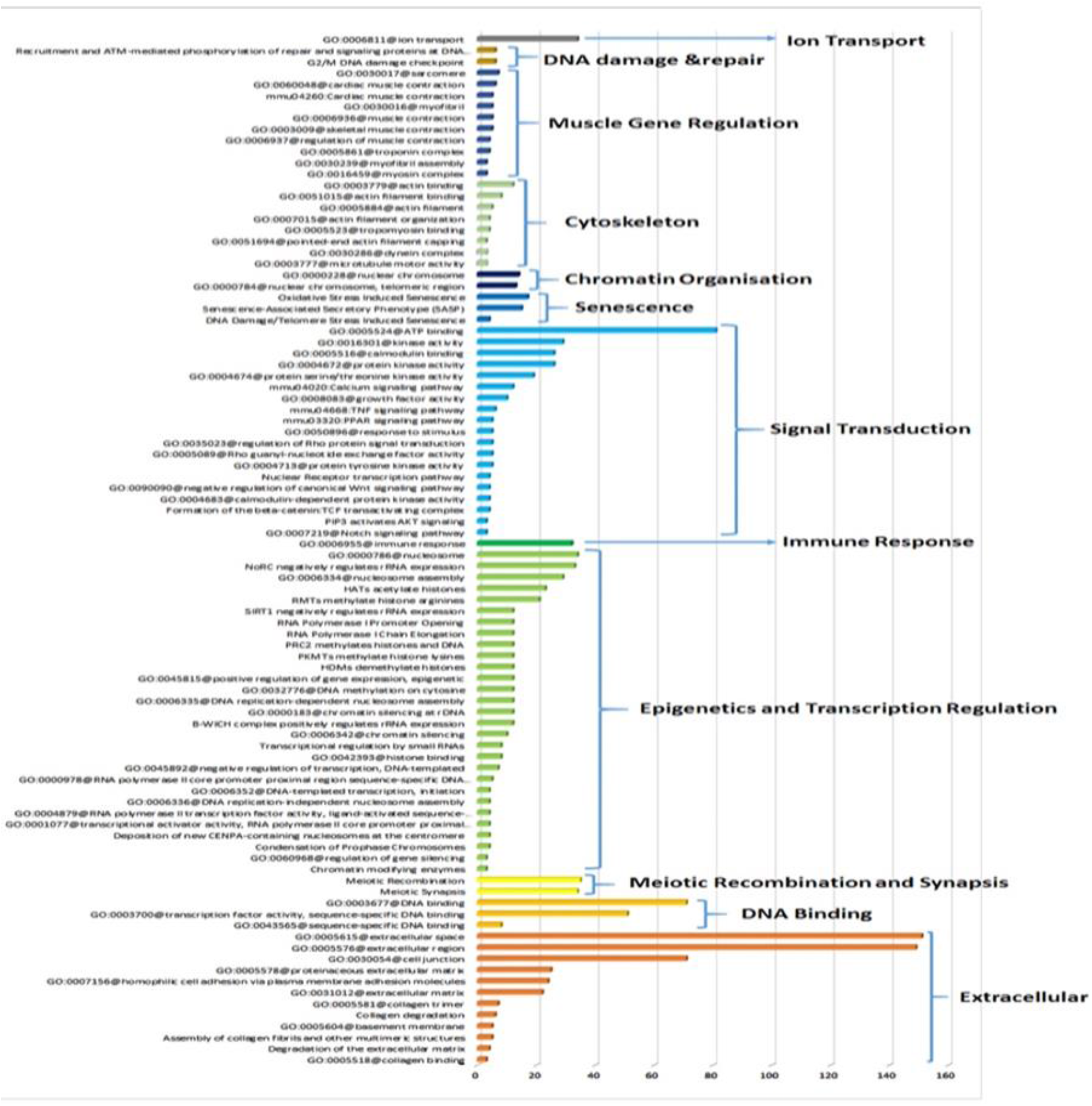
Majorly dysregulated pathways in mutants irrespective of surface conditions.

**Figure Supplementary 3:**
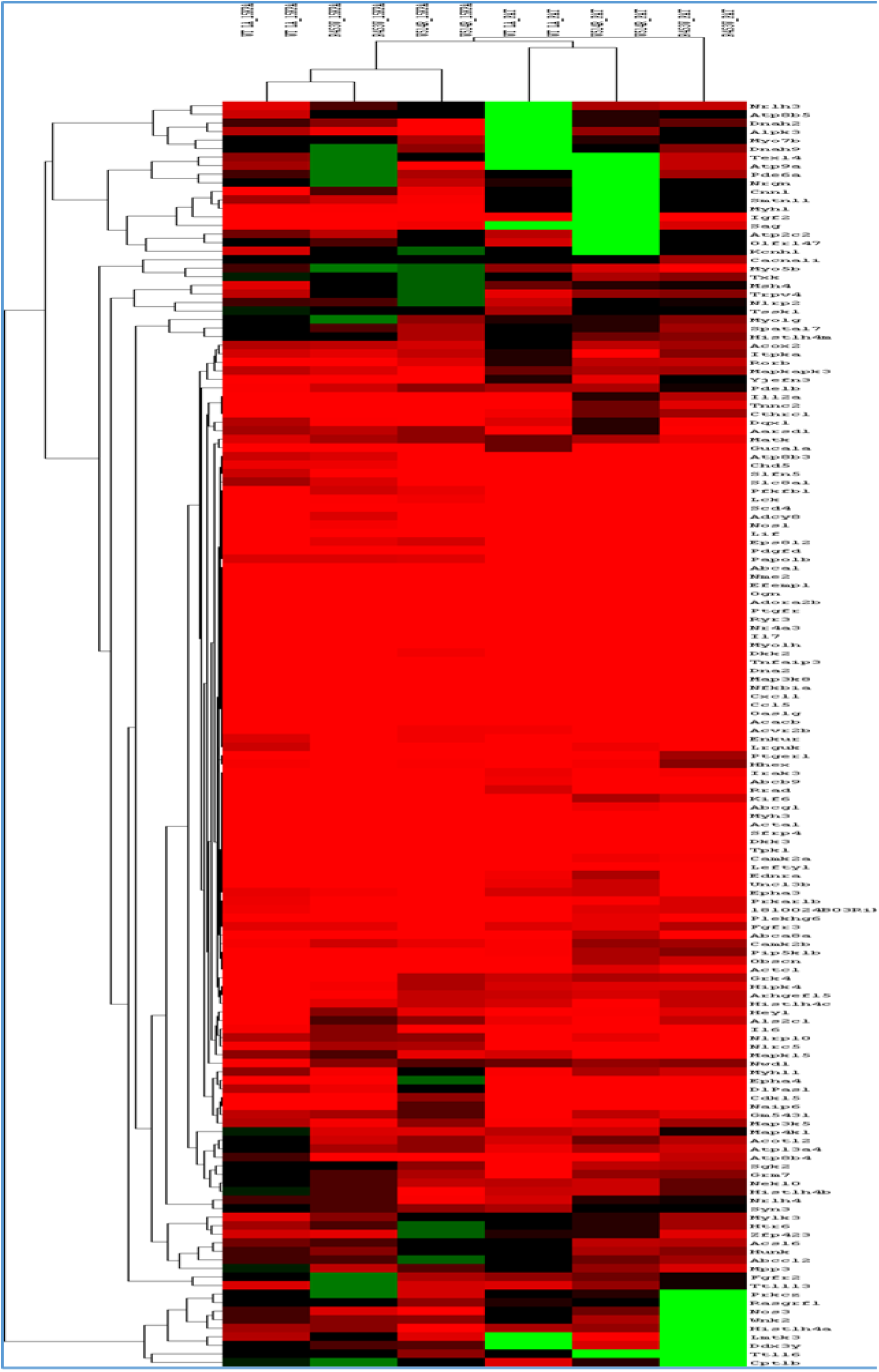
Heatmap of signaling pathways in 6 samples.

